# Genetic, bioacoustic and morphological analyses reveal cryptic speciation in the warbling vireo complex (*Vireo gilvus*: Vireonidae: Passeriformes)

**DOI:** 10.1101/2021.07.12.452121

**Authors:** AM Carpenter, BA Graham, GM Spellman, J Klicka, TM Burg

## Abstract

Cryptic species are closely related taxa that are difficult to separate morphologically, but are reproductively isolated. Here we examine the warbling vireo complex (*Vireo gilvus*), a widespread songbird speculated to be comprised of more than one cryptic species. We included three taxa within the complex: two of the western (*Vireo gilvus swainsonii* and *V. g. brewsteri*) subspecies and the single eastern (*V. g. gilvus*) subspecies. We used mtDNA and microsatellite loci to assess the congruence of genetic data to the current subspecies boundaries. We then incorporated bioacoustic, morphometric, and ecological niche modeling analyses to further examine differences. We found two genetic groups with mtDNA analysis. Microsatellite analyses revealed four genetic groups: an eastern group, a Black Hills group and two western groups that do not agree with current western subspecies boundaries based on phenotypic data. Our results suggest that eastern and western warbling vireos have been reproductively isolated for a long period of time and therefore, may be best treated as separate species; however, more research into areas of contact to examine the presence of hybridization is advised before making a taxonomic revision. Differences between the two western genetic groups appear less clear, requiring additional research.

## INTRODUCTION

Under the biological species concept, “species” are defined as groups of natural interbreeding populations that are reproductively isolated from other such groups (Mayr, 1942). The process of speciation occurs when reproductive isolating barriers accrue to the extent that gene flow between closely related populations greatly diminishes or becomes absent altogether (Coyne & Orr, 2004). Isolation is caused by numerous factors, including behavioral (mating displays), temporal (timing of migration or breeding), and habitat differences (Irwin, 2000; Delmore *et al*., 2016; Cristescu *et al*., 2012). Our understanding of the various forces driving speciation is important in identifying how biodiversity is generated, particularly in the case of cryptic speciation.

Cryptic species were first recognized by Derham (1718) and later defined as closely related taxa that are difficult to separate morphologically, but are reproductively isolated from one another (Lincoln *et al*., 1998; Winker, 2005). Such subtleties in morphological characters between populations leads to subspecies designations within a single species complex, yet the significance of this taxonomic category has been questioned (Ball & Avise, 1992; Zink, 2004). One result is underestimating biological diversity, as one widespread species complex could be in fact multiple, geographically restricted species (Eme *et al*., 2017). Research delimiting species boundaries has become very active over the past decade with advances in technology (Flot, 2015). Specifically, the prevalence of molecular techniques has aided in the identification of cryptic species across diverse taxa (Delić *et al*., 2017; Silva *et al*., 2018), including birds (Isler & Whitney, 2011; Irwin *et al*., 2011). Such genetic studies have played a crucial role in ascertaining phylogenetic relationships and gene flow between subspecies populations, factors that are of consideration when elevating groups to species status which has both biological (Toews & Irwin, 2008) and conservation (Mace, 2004; Murphy *et al*., 2011) importance. Other explanatory variables including acoustic structure (Irwin *et al*., 2008; Toews & Irwin, 2008), ecological niche modeling (Berzaghi *et al*., 2018), and morphological data (Alström *et al*., 2011) have been incorporated to further clarify the taxonomic status of cryptic avian species.

The genus *Vireo* (Vieillot, 1808) (Order: Passeriformes; Family: Vireonidae) contains 34 species and is found throughout the New World (Gill & Donsker, 2019). Thirteen species breed in North America, occupying a mosaic of habitats including temperate and boreal forests, shrublands, and woodland edges (Gill & Donsker, 2019; Mejías *et al*., 2020). Reduction of morphological divergence and similar plumage patterns have been recognized among most species (Hamilton, 1962), postulating that ecological differences could have been a mode of speciation in *Vireo* (Hamilton, 1959; 1962; Barlow 1980). Additionally, a recent study found phylogenetic history strongly shapes song structure and morphology among vireos (Mejías *et al*., 2020). Several cryptic species have been rectified from species complexes within *Vireo*, including Hutton’s vireo, *Vireo huttoni* (Cassin, 1851) (Cicero & Johnson, 1992), blue-headed vireo, *Vireo solitarius* (Wilson, 1810) (Johnson, 1995; Cicero & Johnson, 1998), and red-eyed vireo, *Vireo olivaceus* (Linnaeus, 1766) (Slager *et al*., 2014; Battey & Klicka, 2017). The discovery of these previously unrecognized cryptic species suggests *Vireo* may contain additional cryptic species, so further scrutiny of other complexes within the genus is suggested (Johnson, 1995).

The warbling vireo (*Vireo gilvus* Vieillot 1808) complex taxonomy follows subtle geographic variation of morphological and plumage characters to recognize five subspecies (Phillips, 1991) divided into two groups (AOU, 1998): a western *swainsonii* group containing the subspecies *Vireo gilvus swainsonii* (Baird, 1858), *Vireo gilvus brewsteri* (Ridgway, 1903), *Vireo gilvus sympatricus* (Phillips, 1991), and *Vireo gilvus victoriae* (Sibley, 1940), and an eastern group *gilvus* with the single subspecies *Vireo gilvus gilvus* (Vieillot, 1808). Two of the western subspecies, *V. g. sympatricus* and *V. g. victoriae*, are endemic to regions of Mexico, which fall out of the scope of this study. The remaining three, two of the western (*swainsonii* and *brewsteri*) and the eastern (*gilvus*), subspecies (referred to herein by their subspecies epithet) are Neotropical-Nearctic migrants that breed throughout large parts of Canada and the United States, with sympatric distributions in western North America (Fig. 1) (Phillips, 1991; AOU, 1998). The eastern subspecies is diagnostic from these two western subspecies by its larger size, longer (7.5-mm) and deeper bills (3.6-4.1 mm) with a paler upper mandible, brighter plumage, and p10 is longer vs the primary covert feathers (by 2-8 mm) in western warbling vireos only (Phillips, 1991; Pyle, 1997). Between the two western subspecies, diagnoseable characters are more overlapping; *swainsonii* is a smaller bird than *brewsteri*, but both of their plumages are dull with variations of olive and gray color, and wing and bill morphologies are similar (Phillips, 1991; Pyle, 1997). It has been postulated that *gilvus* and *swainsonii* groups are two separate species (Sibley & Monroe, 1990; AOU, 1998), but more study across the range is sought after (Gardali & Ballard, 2020). Limited analyses of mitochondrial DNA (mtDNA) show divergence of 2.6% for ND2 (Slager et al., 2014), 3.0 – 3.5% for cytochrome b (Murray *et al*., 1994; Lovell *et al*., 2021), and ≥ 3.7% for COI (Hebert *et al*., 2004). Differences in morphology (Lovell, 2010), song (Howes-Jones, 1985; Lovell, 2010), habitat (Semenchuk, 1992; Lovell *et al*., 2021), molt and migratory behavior (Voelker & Rohwer, 1998), and nuclear DNA (Lovell *et al*., 2021) have also been discovered. The applicability of some of these results has been queried, however, as limitations of these previous studies include small sample sizes and restricted sampling of geographical areas and subspecies (e.g., *swainsonii* and *gilvus*) (Gardali & Ballard, 2020).

**Figure 1.**
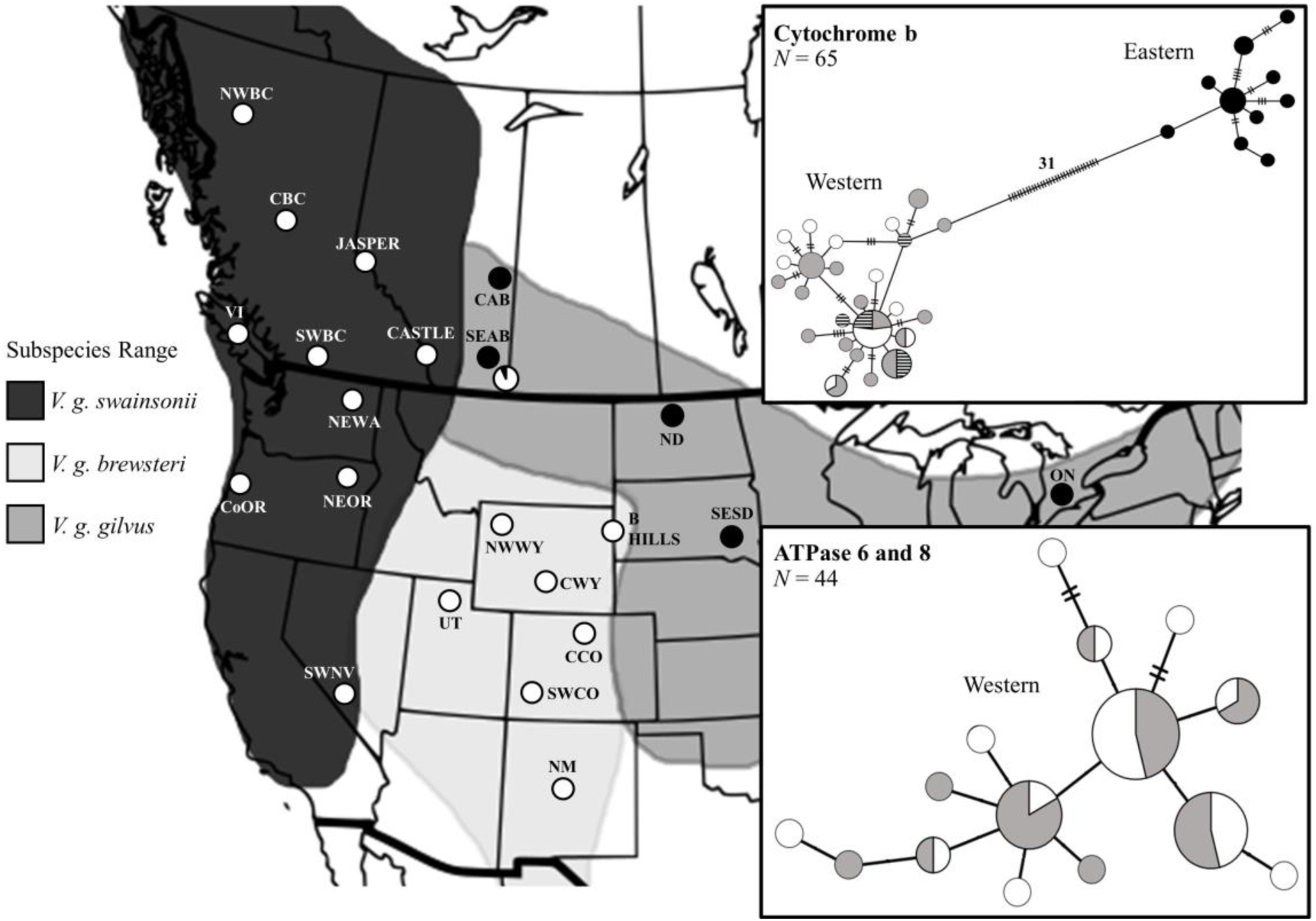
Range distributions of the three warbling vireo subspecies in this study based on Phillips (1991) and AOU (1998). Circles on the map show sampling locations included in the *cyt b* analysis (top inset). Black circles are eastern haplotypes, whereas white circles are western haplotypes. The eastern (Medicine Hat; M HAT) and western (Cypress Hills; C HILLS) groups from the SEAB population are shown separately. The ATPase 6 and 8 results are not on the map (bottom inset) and do not include the eastern group. Colors in the haplotype networks correspond to our microsatellite genetic groups: eastern (black), north-western (grey), south-western (white), and the Black Hills (striped). Each circle is a haplotype, and its size is proportional to how many individuals share that haplotype. Cross hatches in the haplotype networks represent more than one nucleotide difference. Refer to Table 1 for population abbreviations and sample sizes.

Our study aims to fill the knowledge gap about the geographic variation in population genetics across the range of the warbling vireo complex. The use of both mitochondrial and nuclear markers allows for the elucidation of evolutionary history and contemporary genetic patterns, avoiding the limitations of single marker inferences. We implement bioacoustic and morphometric analyses in the context of the subspecies designations and genetic groups to determine the levels of variation among them independently. Finally, the range boundaries between genetic groups are visualized using contemporary ecological niche modeling to compare to the subspecies boundaries. This approach can be powerful when investigating the distributions and differences between cryptic species (Rissler & Apodaca, 2007). Specifically, we ask:

1. Are mtDNA and microsatellite genetic groups congruent to the *swainsonii*, *brewsteri*, and *gilvus* subspecies boundaries?
2. Are there differences in acoustic and morphological characters among eastern and western subspecies? Among eastern and western genetic groups? And
3. Are there differences in acoustic and morphological characters between the western subspecies? Between the western genetic groups?

## METHODS

### Sample Collection

From 2007 – 2019, we collected 194 warbling vireo samples from adult birds in 18 populations. Birds were sampled during the breeding season, end of May through to early July, to avoid migrants and reduce the capture of family groups. Each bird was caught by mist net using song playbacks, banded, and a small blood sample (< 50 µL) was taken from the brachial vein for genetic analyses. Four morphological measurements were collected from each bird using calipers to 0.1 mm (bill length, bill depth, bill width, and tarsus length), wing chord was measured using a wing ruler to the nearest millimeter, and birds were weighed in grams (g) to one decimal point. Additionally, we obtained 184 blood or tissue samples, and corresponding morphological data, if available, from museum collections (Table 1). All blood and tissue samples were stored in 95% ethanol. DNA was extracted using a modified Chelex procedure (Walsh *et al*., 1991; Burg & Croxall, 2001).

**Table 1.** Population abbreviations and the number of individuals (*N*) used in cytochrome b (*cyt b*), ATPase 6 and 8 (*ATP*), microsatellite (msat), and morphometric analyses. Populations with an asterisk include some museum samples.

| **Population ID** | ***cyt b*** | ***ATP*** | ***msat*** | **morphometric** |
| --- | --- | --- | --- | --- |
| NWBC (North-west British Columbia) | 3 | 5 | 4 | 6 |
| CBC (Central British Columbia) | 2 | 4 | 24 | 24 |
| JASPER (Jasper National Park, Alberta) | 2 | 2 | 13 | 10 |
| BANFF (Banff National Park, Alberta) | - | - | 15 | 11 |
| CASTLE (Castle Provincial Park, Alberta) | 1 | 8 | 17 | 9 |
| WLNP (Waterton Lakes National Park, Alberta) | - | 4 | 9 | 8 |
| NEWA (North-east Washington) | 3 | - | 14 | 14 |
| SWBC (South-west British Columbia) | 2 | - | 5 | 5 |
| NWWA (North-west Washington)* | - | - | 1 | - |
| VI (Vancouver Island)* | 1 | - | 4 | - |
| CWA (Central Washington)* | - | - | 20 | - |
| NEOR (North-east Oregon) * | 3 | 3 | 10 | 3 |
| SEOR (South-east Oregon)* | - | - | 1 | - |
| CoOR (Coastal Oregon) | 1 | 2 | 2 | - |
| CoCA (Coastal California)* | - | - | 3 | - |
| NECA (North-east California) | - | - | 3 | 3 |
| SECA (South-east California)* | - | - | 4 | - |
| SWNV (South-west Nevada)* | 1 | - | 10 | - |
| NENV (North-east Nevada)* | - | - | 10 | - |
| UT (Utah) | 1 | 4 | 4 | 4 |
| NM (New Mexico)* | 1 | - | 8 | 1 |
| SWCO (South-west Colorado)* | 1 | 1 | 13 | - |
| CCO (Central Colorado)* | 2 | 3 | 13 | 4 |
| CWY (Central Wyoming)* | 1 | - | 6 | - |
| NWWY (North-west Wyoming)* | 1 | 1 | 10 | 1 |
| SWMT (South-west Montana) | - | 7 | 29 | 26 |
| CMT (Central Montana)* | - | - | 4 | - |
| B HILLS (Black Hills)* | 9 | - | 36 | - |
| SEAB (South-east Alberta) | 22 | - | 24 | 22 |
| CAB (Central Alberta)* | 1 | - | 1 | - |
| SK (Saskatchewan) | - | - | 10 | 11 |
| ND (North Dakota) | 4 | - | 16 | 17 |
| SESD (South-east South Dakota)* | 1 | - | 1 | - |
| MN (Minnesota)* | - | - | 4 | - |
| IL (Illinois)* | - | - | 8 | - |
| OH (Ohio)* | - | - | 5 | - |
| ON (Ontario)* | 2 | - | 17 | - |
| **Total number of samples (*N*)** | **65** | **44** | **378** | **179** |

### DNA Amplification

#### mtDNA

We amplified two mtDNA genes: cytochrome b (*cyt b*) and ATPase 6 and 8. The 915 bp (base pair) cyt b fragment was amplified using the primers H15944 (Lovell *et al*., 2021) and L14990 (Sorenson *et al*., 1999) in 65 warbling vireos from populations of all three subspecies (Table 1). The second gene, a 503 bp fragment of ATPase 6 and 8, was amplified using the primers ATP H534ch (Lait & Burg, 2013) and L8929 COII (Sorenson *et al*., 1999) in 44 warbling vireos from only populations of the two western subspecies, *swainsonii* and *brewsteri*, to further examine their evolutionary histories (Table 1). The 25 µL polymerase chain reaction (PCR) mixture was the same for both mtDNA genes using 5x Green GoTaq® Flexi buffer (Promega), 0.2 mM dNTPs, 2 mM MgCl_2_ (Promega), 0.5 µM of each H and L primer, and 0.5 U GoTaq® Flexi polymerase (Promega). The following thermocycling profile was used to amplify DNA for both mtDNA genes: two minutes at 94°C, 45 seconds at 58°C, one minute at 72°C; 37 cycles of 30 seconds at 94°C, 45 seconds at 58°C, one minute at 72°C, concluding with an extension period of five minutes at 72°C. Sequencing for both *cyt b* and ATPase 6 and 8 genes were completed by NanuQ (McGill University, Montreal, QC, Canada).

#### Microsatellites

A subset of warbling vireo samples (*N* = 6) from geographically distant areas were screened with 57 passerine microsatellite loci. A total of 30 microsatellite loci optimized successfully, 14 of which were polymorphic and used in our study: *BCV2-6*, *BCV4-5*, *BCV4-6New* (Barr *et al*., 2007), *Ck.1B5D*, *Ck.4B6D*, *Ck.2A5A*, *Ck.1B6G* (Tarr & Fleischer, 1998), *Hofi5* (Hawley, 2005), *Pocc1* (Bensch *et al*., 1997), *ApCo46* (Stenzler & Fitzpatrick, 2002), *Lox1* (Piertney *et al*., 1998), *Ase18* (Richardson *et al*., 2000), *Ppi2New* (Martinez *et al*., 1999), and *CmAAAG30* (Williams *et al*., 2004) (see Supporting Information, Table S1). All PCRs were conducted in a 10 µL volume using FroggaBio buffer, 0.2 mM dNTPs, 0.5 µM of forward primer, 1 µM of the reverse primer, 0.05 µM of the M13 fluorescently labeled primer, and 1 U of FroggaBio Taq. The following thermocycling profile was used to amplify DNA: one cycle for one minute at 94°C, 45 seconds at T_1_, and one minute at 72°C; seven cycles of one minute at 94°C, 30 seconds at T_1_, and 45 seconds at 72°C; 25 cycles for 30 seconds at 94°C, 30 seconds at T_2_, and 45 seconds at 72°C; and one cycle for 72°C at five minutes. Five loci (*BCV4-6New*, *Hofi5*, *ApCo46*, *Lox1*, and *Ppi2New*) were amplified using T_1_=50°C and T_2_=52°C, five loci (*BCV2-6*, *BCV4-5*, *Ck.1B5D*, *Ck.2A5A*, and *Ck.4B6D*) were amplified using T_1_=52°C and T_2_=54°C, and four loci (*Pocc1*, *Ase18*, *CmAAAG30*, and *Ck.1B6G*) were amplified using T_1_=55°C and T_2_=57°C. For three loci (*BCV4-5*, *ApCo46*, and *Ppi2New*) the second step was increased from 25 to 31 cycles. PCR products were run on a 6% acrylamide gel on a LI-COR 4300 DNA Analyzer (LI-COR Inc., Lincoln, NE, USA). To maintain consistency of scoring alleles, the same three positive controls of known sizes were amplified and run with each load for each locus. All alleles were scored manually by visual inspection and checked independently by a second person. Samples from each load was re-run at a later date to ensure the accuracy of allele scores.

### Genetic Analyses

#### MtDNA

Sequences were checked and aligned using MEGA X 10.1 (Kumar *et al*., 2018). *Cyt b* sequences were compared to a reference warbling vireo sequence from GenBank (AY030111). We constructed minimum spanning haplotype networks from variable sites at both genes to examine historic genetic differences among subspecies. Minimum spanning haplotype networks were created for ATPase 6 and 8 and *cyt b* using default settings in Population Analysis with Reticulate Trees 1.7 (PopART) software (Bandelt *et al*., 1999).

We estimated coalescence times between the eastern and western genetic groups, between the two western genetic groups, and between these genetic groups and the Black Hills group (see results) by applying a molecular clock of 2% per million years (Brown *et al*., 1979) to our *cyt b* data, a rate used in Passeriformes (Weir & Schluter, 2008). Additionally, we estimated a second coalescence time between the two western genetic groups using data from ATPase 6 and 8 with the same 2%/Myr rate (Brown *et al*., 1979).

#### Microsatellites

A total of 378 individuals from 37 populations were included in microsatellite analyses (Table 1). Expected (*H_E_*) and observed (*H_O_*) heterozygosities and the average number of alleles per locus were calculated in GenAlEx 6.5 (Peakall & Smouse, 2006; 2012) (Supporting Information, Table S1). Deviations from Hardy-Weinberg equilibrium and linkage disequilibrium were tested with Genepop 4.7 (Raymond & Rousset, 1995; Rousset, 2008).

Pairwise F_ST_ values were calculated in GenoDive (Meirmans & van Tienderen, 2004) to examine genetic differentiation among populations with more than five individuals. P-values were corrected *post hoc* with the Benjamini-Hochberg (B-H) method to account for multiple comparisons and avoid type I errors (Benjamini & Hochberg, 1995). We chose the B-H method because Bonferroni corrections have been suggested as too severe, often underestimating the number of significant comparisons (Garamszegi, 2006).

We used the program STRUCTURE 2.3.4 (Pritchard *et al*., 2000; Hubisz *et al*., 2009) to examine the number of genetic groups present in the warbling vireo complex using the following settings: admixture model with correlated allele frequencies, sampling locations set as *locpriors*, and a burn-in of 50 000 followed by 100 000 Monte Carlo Markov Chains (McMC) for ten iterations for each K from 1 to 5. We calculated ΔK (Evanno *et al*., 2005) in Structure Harvester (Earl & von Holdt, 2012) and examined posterior probabilities with Bayes rule (Pritchard *et al*., 2000) to determine the optimal K. We tested for the presence of hierarchal structure by running subsequent analyses for each genetic group identified in the initial STRUCTURE runs. Final outputs for each optimal value of K were averaged among the ten iterations and represented graphically using Structure Plot 2.0 (Ramasamy *et al*., 2014).

### Bioacoustic Analyses

To explore acoustic differences, we compared songs from the Xeno-canto repository (www.xeno-canto.org) for three out of the four microsatellite genetic groups (excluding the Black Hills group). Of the 336 song recordings originally downloaded, we measured songs from 47 audio files. When deciding which audio files to measure, we looked for songs with a high signal-to-noise ratio and no overlap from other species; we chose to examine one file for each site to avoid measuring songs from the same bird, thereby avoiding pseudoreplication. For each song, we checked the location where the song was recorded and then assigned it to a genetic group based on our microsatellite genotyping results. Of the 47 audio files 13 were north-western, 26 were south-western, and eight were eastern. From each audio file, up to seven songs were analyzed for five fine-scale measurements: song length (s), minimum frequency (Hz), maximum frequency (Hz), peak frequency (Hz; the frequency of the song with the maximum power), and the syllable delivery rate of each song (syllables/second). To calculate syllable delivery rate, we counted the average number of syllables in each song and then divided this number by the average song length. All values were tested for intercorrelations and, although we originally measured bandwidth (Hz) for each song, this measurement was highly correlated (r = 0.95) with maximum frequency and removed from our analyses. All sound files were converted to WAVE files, as WAVE files can produce different results with respect to fine-scale measurements (Araya-Salas *et al*., 2017). All measurements were done in RavenPro 1.4 (Cornell lab of Ornithology, Ithaca, New York, USA) using the following settings: 50% Hahn window and 256 kHz sampling with 16-bit accuracy. For audio files where more than one song was measured, we averaged all of the songs to produce a single data point for that audio file.

To determine whether the songs of the northwestern, southwestern, and eastern genetic groups are acoustically different based on these five fine-scale measurements, we carried out a discriminate function analysis (DFA). We used the leave-one out classification analysis and reported the percentage of songs that were correctly assigned to each group. We used this same DFA to examine whether there were also acoustic differences among the subspecies. Songs were measured using the same settings as mentioned above for the genetic groups, but each song was assigned to a subspecies based on the information associated with the audio file. In those instances where a subspecies rank was not included with the file (*N* = 22), we assigned it to a subspecies based on the geographic location of where the audio file was recorded in respect to the subspecies boundaries as described by Phillips (1991). Of the 47 audio files 24 were *swainsonii*, 11 were *brewsteri*, and 12 were *gilvus*. DFAs were performed in SPSS 23.0 (SPSS Inc., Chicago, IL, USA).

Lastly, songs were compared among the subspecies and among the genetic groups independently using a multivariate analysis of variance (MANOVA) in SPSS 23.0 (SPSS Inc., Chicago, IL, USA). The first and second canonical axes were used our dependent variables and either genetic groups or subspecies was used as our independent variable. A Tukey’s *post hoc* test was done to evaluate whether songs were different among the subspecies and among the genetic groups.

### Morphometric Analyses

Using the genetic groups from our microsatellite data, we did a MANOVA in SPSS 23.0 to test for morphological differences between them. Six morphological measurements were compared: mass, wing chord, bill length (from nares to tip), bill depth, bill width (posterior end of nostrils), and tarsus length among 179 individuals assigned to a genetic group (88 north-western, 56 south-western, 35 eastern) (excluding the Black Hills group) (Table 1).We included the genetic groups as our dependent variable, and latitude and bird bander as covariate variables in our model due to the potential of bird bander biases and effects of latitude on morphology. These same 179 individuals were also assigned to one of the three subspecies (108 *swainsonii*, 36 *brewsteri*, and 35 *gilvus*) to compare morphological differences. For this analysis, we used subspecies as our dependent variable and again used latitude and bird bander as covariates in our model. A least-squared difference test was used to examine the significance of morphological differences among the genetic groups and among the subspecies independently.

Similar to our methodology of assessing acoustic differences, we also performed a DFA on morphological traits. The six morphological characters were used as our dependent variables, and we used the leave-one out classification analysis to report the percentage of individuals that were correctly assigned to each genetic group and subspecies based on morphology. The use of DFA to diagnose vocal and morphological differences between subspecies follows techniques outlined in Demko *et al*. (2020).

### Ecological Niche Modelling

We ran ecological niche models for three of the four genetic groups (Black Hills was not included) to project each of their ranges and explore the role of climate variables involved in their distributions. Using Maxent 3.4.1 (Phillips *et al*., 2006) with WorldClim BioClim rasters (Hijmans *et al*., 2005), predictive models output estimates of suitable areas of habitat for a species based on the environmental conditions from occurrence datasets of that species (Elith & Leathwick, 2009). Occurrence records were comprised of sampling locations from this study (*N* = 105) and eBird datasets (www.ebird.org; Cornell Lab of Ornithology, 2019) (*N* = 4593). To restrict occurrences to breeding populations, only observations from late May to early August were included and occurrences points were assigned to a genetic group based on our microsatellite data. Wallace 1.0.6 in R (Kass *et al*., 2017) was used to spatially thin occurrence points by 10 km to avoid over representation of a given area. Final occurrence sets for each genetic group was north-western group *N* = 590, south-western group *N* = 718, and eastern group *N* = 1114.

Nineteen variables at 2.5 arcmin resolution were downloaded from the WorldClim BioClim dataset and tested for correlations. Once we removed all but one variable with strong correlations (> 0.7), we were left with 8 – 12 variables depending on each genetic group’s model; BIO4 and BIO19 were not used in any model. Ten replicates were run for each genetic group’s occurrence points and specific set of BioClim rasters using the 1.0 regularization multiplier with auto features, 10000 background points, and a cross-validation method in Maxent.

## RESULTS

### MtDNA

A 915 bp fragment of *cyt b* was successfully amplified in 65 warbling vireos from 22 populations across all three subspecies ranges (GenBank accession numbers MZ020223 - MZ020287). We identified two mtDNA lineages, separating eastern and western groups by 31 fixed nucleotide differences (Fig. 1). One population, SEAB, contained birds with haplotypes from both groups (six of the 22 individuals had eastern haplotypes). Overall, we identified 35 haplotypes, of which ten haplotypes were exclusive to the eastern lineage. Average sequence divergence for *cyt b* between the eastern genetic group and both western genetic groups and the Black Hills was 4.0% (± 0.1), between the two western genetic groups was 0.4% (± 0.01), and between each of the two western genetic groups and the Black Hills was 0.3% (± 0.01). Applying a 2%/Myr molecular clock places an estimated coalescence time of 2 Mya between the eastern group and the two western groups and the Black Hills, 0.2 Mya between the two western groups, and 0.15 Mya between the two western groups and the Black Hills.

ATPase 6 and 8 was successfully amplified in 44 warbling vireos from 11 populations across the two western subspecies ranges (GenBank accession numbers MZ020288 - MZ020331). We found a single mtDNA lineage with no clear partitioning between subspecies (Fig. 1). We identified 15 haplotypes, one of which is shared by 13 of the 44 individuals (30%). Average sequence divergence was 0.3% (± 0.01) between the two western genetic groups: north-western, formerly *swainsonii*, (*N* = 24) and south-western, formerly *brewsteri*, (*N* = 21). Applying a 2%/Myr molecular clock, this places an estimated coalescence time of 150,000 years between them.

### Microsatellites

Observed and expected heterozygosities were comparable across thirteen of the fourteen loci. *BCV4-6New* had lower *H_O_* (0.59) than *H_E_* (0.71), which could be attributed to an excess of homozygotes genotyped at this locus. The average number of alleles per locus ranged from 2 to 15 (Supporting Information, Table S1). We found no significant departures from Hardy-Weinberg equilibrium or linkage disequilibrium at any of the 14 microsatellite loci.

STRUCTURE identified two genetic groups on the initial run supported by Bayes rule (0.11) and ΔK (−13458). We found a distinct west – east split along the Great Plains dividing western populations from all eastern populations, with the exception of central Montana (CMT) and southeast Alberta (SEAB) (Fig. 2A; Fig. 3). Within the SEAB population, seven out of 24 individuals assigned to the eastern genetic group. All eastern populations, and the seven eastern individuals within the SEAB population, were removed to test for additional substructure among western populations independently.

**Figure 2.**
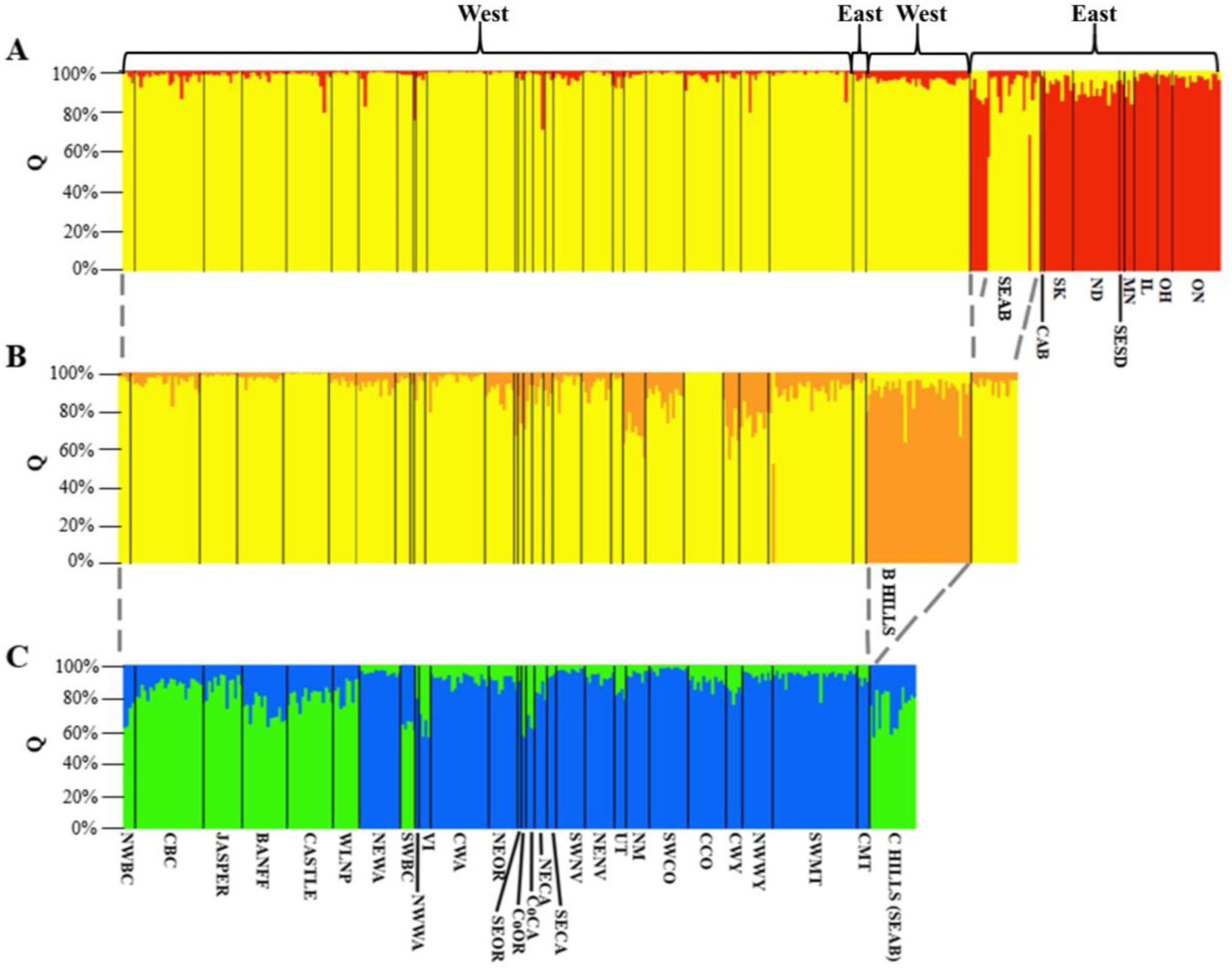
Hierarchical STRUCTURE plots (K = 2) based on genotypes from 14 microsatellite loci. Each bar represents a single individual, and the Q value is the percent ancestry of that individual to each genetic group. Populations with additional hierarchical structure are shown in yellow. **(A)** All populations (red = eastern group), **(B)** Western populations and the Black Hills (orange), **(C)** Remaining western populations (north-western = green; south-western = blue). Refer to Table 1 for population abbreviations and sample sizes.

**Figure 3.**
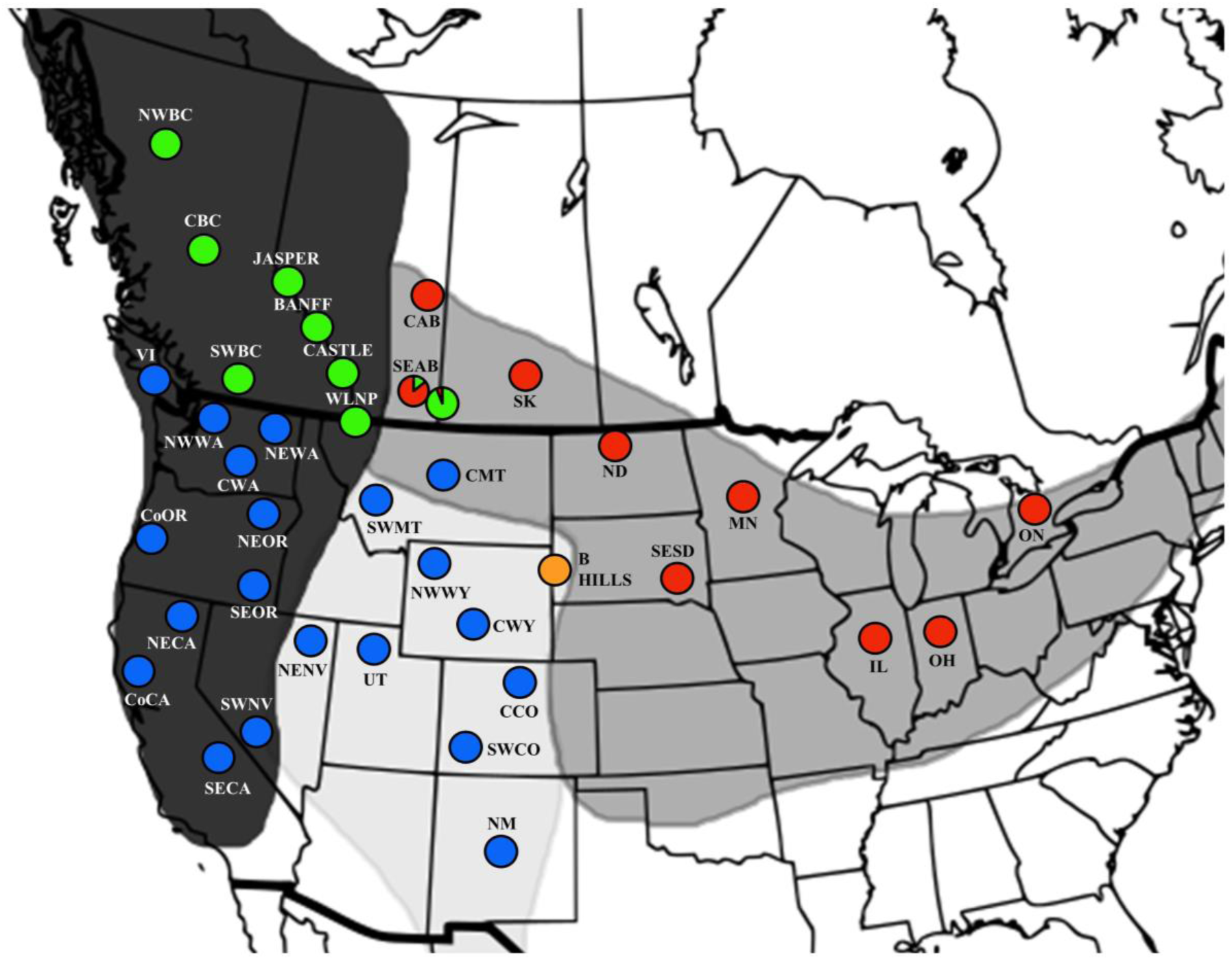
Thirty-seven warbling vireo populations genotyped at 14 microsatellite loci. Colors correspond to the four genetic groups from STRUCTURE: eastern (red), north-western (green), south-western (blue), and the Black Hills (orange) (see Figure 2). The eastern (Medicine Hat; M HAT) and north-western (Cypress Hills; C HILLS) groups from the SEAB population are shown separately. Refer to Table 1 for population abbreviations and sample sizes.

Among the western populations, K = 2 separated all western populations from the Black Hills (Bayes rule = 0.27; ΔK = -11021) (Fig. 2B). A subsequent run was executed after removing the Black Hills population, with two additional genetic groups detected among the remaining western populations (Bayes rule = 1.0; ΔK = 9785). The majority of individuals from each population were assigned Q ≥ 60% to a specific western genetic group, but do not follow the current western subspecies range distributions. For this reason, we will refer to *swainsonii* as the north-western genetic group and *brewsteri* as the south-western genetic group. The north-western group contains seven populations from the *swainsonii* range (NWBC, CBC, JASPER, BANFF, CASTLE, WLNP, SWBC), in addition to one population from the *gilvus* range (SEAB), while the south-western group contains eleven populations from the *swainsonii* range (NEWA, NWWA, VI, CWA, NEOR, SEOR, CoOR, CoCA, NECA, SECA, SWNV), eight populations from the *brewsteri* range (NENV, UT, NM, SWCO, CCO, CWY, NWWY, SWMT), and one population from the *gilvus* range (CMT) (See Table 1 for abbreviation names; Fig. 2C). Altogether, STRUCTURE identified four genetic clusters: an eastern group, two western groups (north-western and south-western), and the Black Hills (Fig. 3).

Based on the microsatellite genetic groups, we split the southeast Alberta (SEAB) population into two separate populations, Cypress Hills (C HILLS) and Medicine Hat (M HAT) for the pairwise F_ST_ analysis. We found high levels of population genetic structure for warbling vireos based on pairwise F_ST_ comparisons. Of the 23 populations examined, 152 of the 253 comparisons were significant (Table 2). Ninety of the significant comparisons separated the eastern group from both the north-western and south-western groups (*P* ≤ 0.01). Three south-western populations, north-east Oregon (NEOR), south-west Nevada (SWNV), and New Mexico (NM), were significantly different from all five northwestern populations (*P* ≤ 0.01) in British Columbia and Alberta: central British Columbia (CBC), Jasper National Park (JASPER), Banff National Park (BANFF), Castle Provincial Park (CASTLE), Waterton Lakes National Park (WLNP). Overall, pairwise F_ST_ comparisons found little genetic differentiation within conspecific populations.

**Table 2.** Pairwise F_ST_ comparisons for 23 populations (*N* > 5) based on 14 microsatellite loci genotypes. F_ST_ values are below the diagonal line, with corresponding P-values above the diagonal line and significant values bolded.

|  | CBC | JASPER | BANFF | CASTLE | WLNP | NEWA | CWA | NEOR | SWNV | NENV | NM | SWCO | CCO | CWY | NWWY | SWMT | B HILLS | C HILLS | M HAT | SK | ND | IL | ON |
| --- | --- | --- | --- | --- | --- | --- | --- | --- | --- | --- | --- | --- | --- | --- | --- | --- | --- | --- | --- | --- | --- | --- | --- |
| CBC | - | 0.29 | **0.02** | 0.06 | 0.18 | 0.17 | **<0.01** | **<0.01** | **<0.01** | 0.07 | **<0.01** | **<0.01** | 0.21 | 0.13 | 0.06 | **<0.01** | **<0.01** | **<0.01** | **<0.01** | **<0.01** | **<0.01** | **<0.01** | **<0.01** |
| JASPER | 0.00 | - | **0.03** | 0.26 | 0.04 | 0.13 | **0.01** | **<0.01** | **0.01** | 0.04 | **<0.01** | **0.01** | 0.16 | 0.12 | 0.06 | 0.06 | **<0.01** | 0.06 | **<0.01** | **<0.01** | **<0.01** | **<0.01** | **<0.01** |
| BANFF | 0.02 | 0.02 | - | 0.13 | 0.60 | 0.43 | **0.01** | **0.01** | **<0.01** | 0.04 | **0.01** | **0.03** | 0.15 | 0.28 | 0.04 | **0.03** | **<0.01** | **<0.01** | **<0.01** | **<0.01** | **<0.01** | **<0.01** | **<0.01** |
| CASTLE | 0.01 | 0.01 | 0.01 | - | 0.27 | 0.05 | 0.12 | **<0.01** | **<0.01** | 0.37 | **0.01** | 0.15 | 0.50 | 0.37 | 0.41 | 0.13 | **<0.01** | **0.03** | **<0.01** | **<0.01** | **<0.01** | **<0.01** | **<0.01** |
| WLNP | 0.01 | 0.02 | 0.00 | 0.01 | - | 0.18 | **0.01** | **0.01** | **<0.01** | 0.08 | **<0.01** | **<0.01** | 0.20 | 0.12 | 0.21 | **0.01** | **<0.01** | **<0.01** | **<0.01** | **<0.01** | **<0.01** | **<0.01** | **<0.01** |
| NEWA | 0.01 | 0.01 | 0.00 | 0.02 | 0.01 | - | 0.43 | 0.13 | 0.64 | 0.60 | **0.02** | 0.39 | 0.70 | 0.38 | 0.19 | 0.44 | **<0.01** | **<0.01** | **<0.01** | **<0.01** | **<0.01** | **<0.01** | **<0.01** |
| CWA | 0.02 | 0.03 | 0.02 | 0.01 | 0.02 | 0.00 | - | 0.45 | **0.03** | 0.76 | **0.01** | 0.70 | 0.49 | 0.76 | 0.30 | **0.03** | **<0.01** | **<0.01** | **<0.01** | **<0.01** | **<0.01** | **<0.01** | **<0.01** |
| NEOR | 0.03 | 0.04 | 0.03 | 0.04 | 0.04 | 0.01 | 0.00 | - | 0.14 | 0.26 | **0.02** | 0.05 | 0.35 | 0.52 | 0.15 | 0.04 | **<0.01** | **<0.01** | **<0.01** | **<0.01** | **<0.01** | **<0.01** | **<0.01** |
| SWNV | 0.05 | 0.04 | 0.05 | 0.04 | 0.05 | 0.00 | 0.02 | 0.02 | - | 0.08 | **0.03** | 0.08 | 0.09 | 0.22 | 0.18 | **<0.01** | **<0.01** | **<0.01** | **<0.01** | **<0.01** | **<0.01** | **<0.01** | **<0.01** |
| NENV | 0.02 | 0.02 | 0.02 | 0.00 | 0.02 | 0.00 | 0.00 | 0.01 | 0.02 | - | 0.15 | 0.73 | 0.78 | 0.68 | 0.85 | 0.37 | **0.03** | 0.09 | **<0.01** | **<0.01** | **<0.01** | **<0.01** | **<0.01** |
| NM | 0.04 | 0.06 | 0.04 | 0.03 | 0.04 | 0.04 | 0.03 | 0.04 | 0.05 | 0.01 | - | 0.27 | 0.32 | 0.78 | 0.20 | 0.06 | 0.22 | **0.02** | **<0.01** | **<0.01** | **<0.01** | **<0.01** | **<0.01** |
| SWCO | 0.03 | 0.03 | 0.02 | 0.01 | 0.03 | 0.00 | 0.00 | 0.02 | 0.02 | 0.00 | 0.01 | - | 0.90 | 0.77 | 0.51 | 0.56 | 0.11 | 0.05 | **<0.01** | **<0.01** | **<0.01** | **<0.01** | **<0.01** |
| CCO | 0.01 | 0.01 | 0.01 | 0.00 | 0.01 | 0.00 | 0.00 | 0.00 | 0.02 | 0.00 | 0.01 | 0.00 | - | 0.89 | 0.25 | 0.83 | **<0.01** | **0.02** | **<0.01** | **<0.01** | **<0.01** | **<0.01** | **<0.01** |
| CWY | 0.02 | 0.02 | 0.01 | 0.00 | 0.02 | 0.00 | 0.00 | 0.00 | 0.02 | 0.00 | 0.00 | 0.00 | 0.00 | - | 0.89 | 0.65 | 0.94 | 0.52 | **<0.01** | **<0.01** | **<0.01** | **<0.01** | **<0.01** |
| NWWY | 0.02 | 0.02 | 0.02 | 0.00 | 0.01 | 0.01 | 0.00 | 0.02 | 0.02 | 0.00 | 0.01 | 0.00 | 0.01 | 0.00 | - | 0.28 | 0.24 | 0.19 | **<0.01** | **<0.01** | **<0.01** | **<0.01** | **<0.01** |
| SWMT | 0.02 | 0.01 | 0.02 | 0.01 | 0.03 | 0.00 | 0.01 | 0.02 | 0.04 | 0.00 | 0.02 | 0.00 | 0.00 | 0.00 | 0.01 | - | **<0.01** | 0.04 | **<0.01** | **<0.01** | **<0.01** | **<0.01** | **<0.01** |
| B HILLS | 0.04 | 0.04 | 0.04 | 0.03 | 0.03 | 0.03 | 0.03 | 0.04 | 0.06 | 0.02 | 0.01 | 0.01 | 0.02 | 0.00 | 0.01 | 0.02 | - | **<0.01** | **<0.01** | **<0.01** | **<0.01** | **<0.01** | **<0.01** |
| C HILLS | 0.03 | 0.02 | 0.04 | 0.02 | 0.04 | 0.05 | 0.03 | 0.04 | 0.05 | 0.02 | 0.03 | 0.02 | 0.02 | 0.00 | 0.01 | 0.02 | 0.02 | - | **<0.01** | **<0.01** | **<0.01** | **<0.01** | **<0.01** |
| M HAT | 0.10 | 0.10 | 0.11 | 0.10 | 0.11 | 0.10 | 0.13 | 0.18 | 0.09 | 0.09 | 0.09 | 0.10 | 0.08 | 0.09 | 0.09 | 0.11 | 0.09 | 0.10 | - | 0.64 | 0.18 | 0.06 | 0.56 |
| SK | 0.12 | 0.10 | 0.13 | 0.10 | 0.14 | 0.12 | 0.13 | 0.17 | 0.10 | 0.11 | 0.09 | 0.10 | 0.08 | 0.08 | 0.08 | 0.10 | 0.08 | 0.09 | 0.00 | - | **0.02** | 0.29 | 0.06 |
| ND | 0.13 | 0.12 | 0.13 | 0.12 | 0.13 | 0.12 | 0.15 | 0.21 | 0.10 | 0.14 | 0.12 | 0.13 | 0.12 | 0.12 | 0.11 | 0.13 | 0.10 | 0.12 | 0.01 | 0.03 | - | **0.01** | **0.03** |
| IL | 0.17 | 0.13 | 0.18 | 0.14 | 0.19 | 0.19 | 0.19 | 0.25 | 0.14 | 0.17 | 0.16 | 0.17 | 0.14 | 0.16 | 0.16 | 0.15 | 0.14 | 0.13 | 0.04 | 0.01 | 0.04 | - | **0.03** |
| ON | 0.16 | 0.16 | 0.18 | 0.16 | 0.19 | 0.16 | 0.19 | 0.25 | 0.14 | 0.16 | 0.14 | 0.14 | 0.13 | 0.15 | 0.14 | 0.15 | 0.13 | 0.14 | 0.00 | 0.02 | 0.02 | 0.03 | - |

### Bioacoustics

For our analysis of song variation in the context of genetic groups, the first canonical function examining syllable delivery rate and song length explained 83% of the observed variance (Wilks’ lambda = 0.55, χ^2^ = 25.3, *P* < 0.001) and songs were significantly distinct among the three genetic groups (Wilks’ lambda = 0.55, F_4,86_ = 5.5, *P* < 0.001) (Fig. 4). Of note are three southwestern birds from Wyoming and Colorado that cluster with the eastern birds. The areas where the songs were recorded is near a contact zone; these could have been recordings from eastern birds and were misidentified. Eastern birds sang longer songs with a higher syllable delivery rate than birds from the two western genetic groups (F_2,44_ = 12.2, *P* < 0.001) (Fig. 5). Songs from the two western genetic groups were not distinct based on the first canonical analysis (*P* = 1). We also found no differences among the three genetic groups for the second canonical function (F_2,44_ = 0.3, *P* = 0.74); 68.1% of the songs were assigned to the correct microsatellite genetic group following cross-validation. Overall, 50% of eastern songs were correctly classified to the eastern genetic group, 84.6% of north-western songs were correctly assigned to the north-western genetic group, and 73.1% of south-western songs were correctly classified to the south-western genetic group.

**Figure 4.**
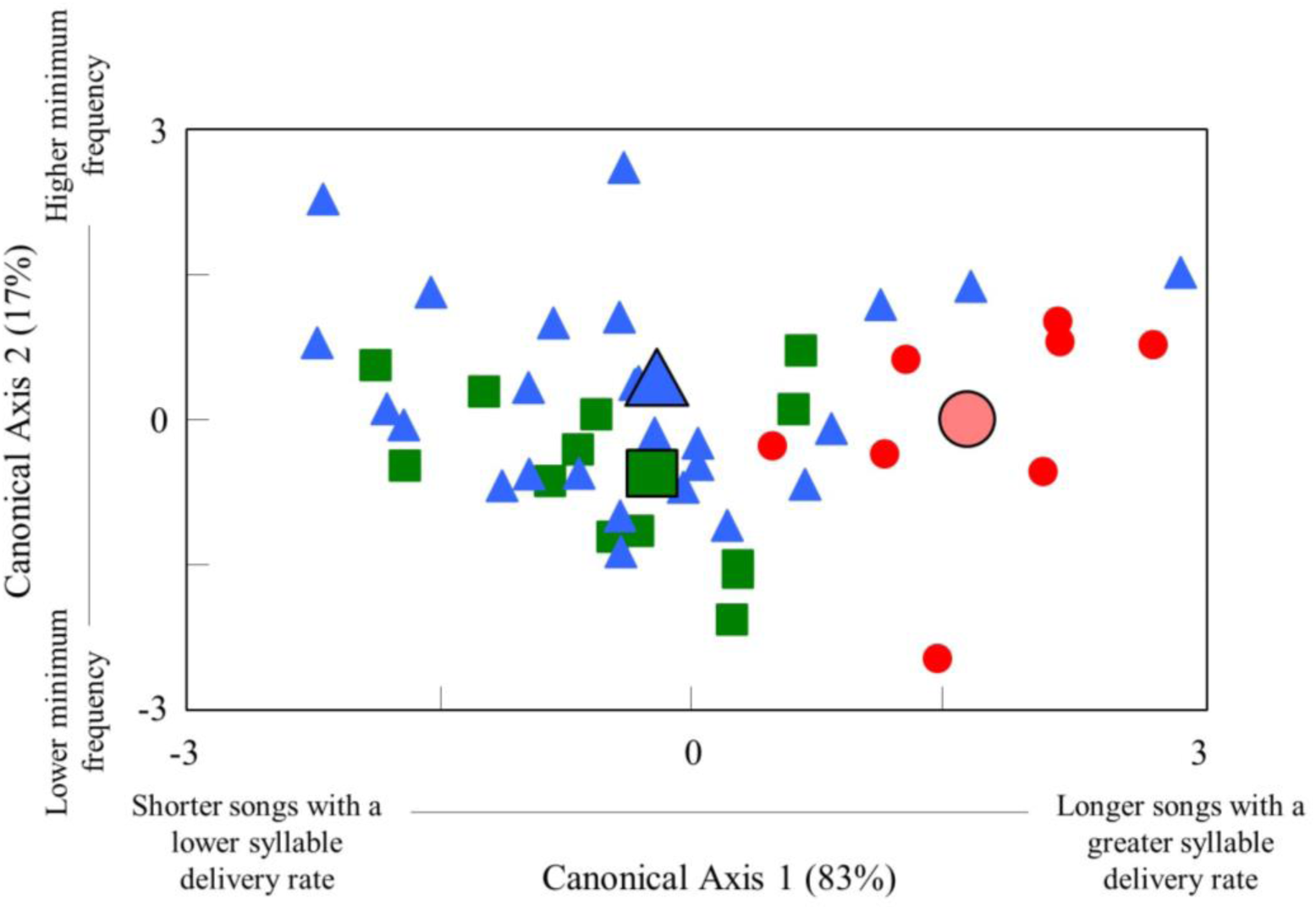
Plots of the first two canonical axes based on discriminate function analysis of warbling vireo songs. Colors and shapes correspond to three of the four microsatellite genetic groups (Black Hills are not included): eastern (red/circle), north-western (green/square), and south-western (blue/triangle). The largest shape represents the mean centroid for each genetic group.

**Figure 5.**
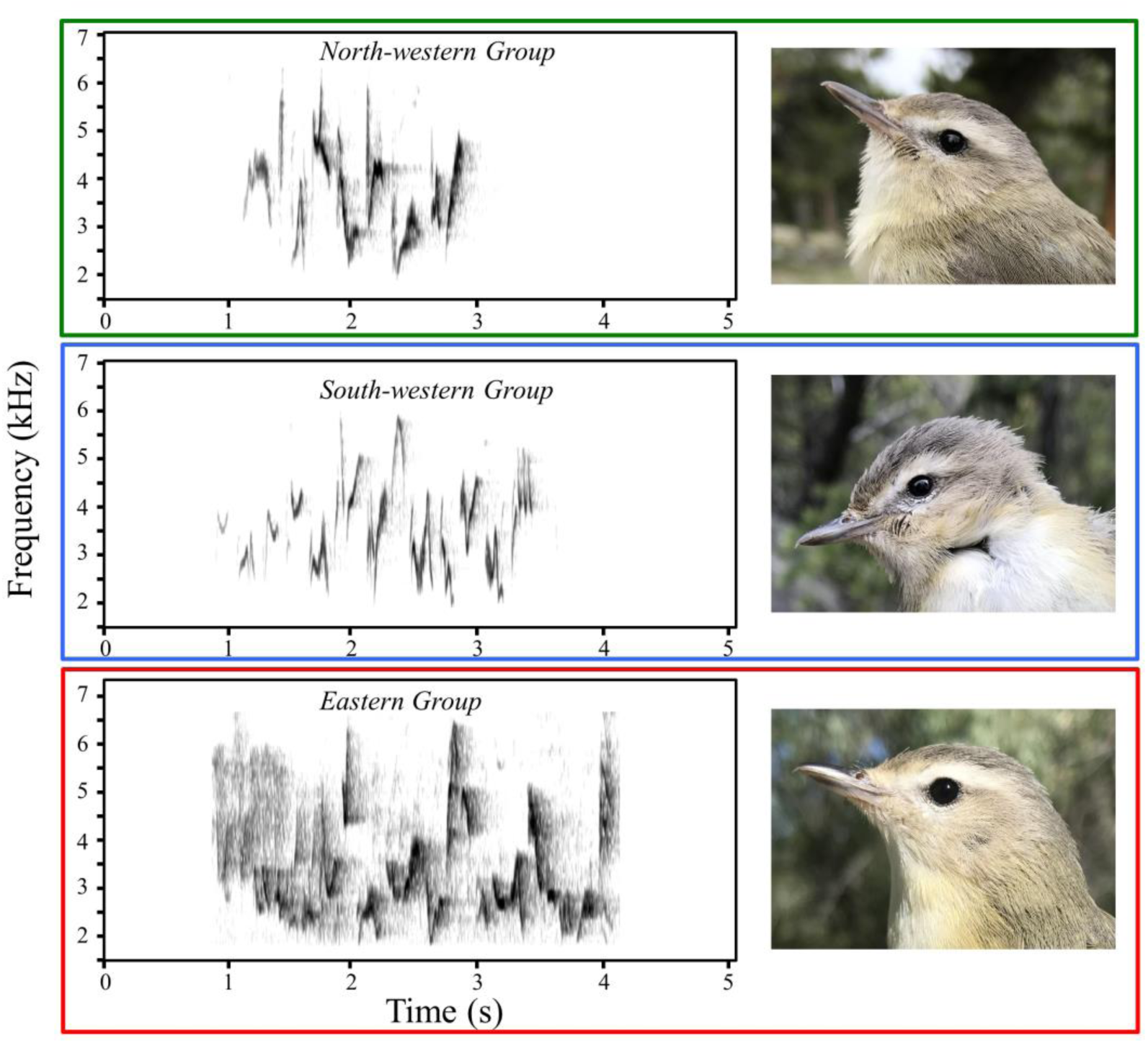
Example song spectrograms and photos for each of the three warbling vireo genetic groups used in our discriminate function analysis of songs. The colors of the outlines of the spectrograms and photos correspond to DFA results in Figure 4. Photos were taken by the first author.

For our analysis using subspecies designations to classify song variation, the first canonical function explained 86% of the observed variance (Wilks’ lambda = 0.50, χ^2^= 29.5, *P* < 0.001). Similar to our analysis using the genetic groups, songs were statistically significant and distinct among subspecies (Wilks’ lambda = 0.5, F_4,86_ = 9.0, *P* < 0.001). Songs from the eastern subspecies are longer and have a higher syllable delivery rate than songs from both of the western subspecies based on the first canonical function (F_2,44_= 17.4, *P* < 0.001). Between the two western subspecies, *swainsonii* songs were shorter and slower paced than *brewsteri* songs (*P* = 0.06). We found no song differences among the subspecies based on the second canonical function (F_2,44_ = 2.8, *P* = 0.08). Overall, 70.2% of the songs were assigned to the correct subspecies following cross-validation. Both *swainsonii* (79.2%) and *gilvus* (72.7%) had high assignment to the correct subspecies, while only 58.3% were correctly assigned as *brewsteri* songs; of note, 25% of *brewsteri* songs were classified as *swainsonii* songs.

### Morphometrics

Morphological differences were found among the three genetic groups (Wilks’ lambda = 0.27, F_12,338_ = 25.8, *P* < 0.001) at five of the six measurements: mass, wing chord, bill depth, bill width, and tarsus; bill length was not significant (see Supporting Information, Table S2). The eastern genetic group is heavier (14.16 g ± 0.19), with longer wings (69.96 mm ± 0.36) and deeper bills (3.96 mm ± 0.05) than north-western (mass: 11.61 g ± 0.13; wing chord: 67.04 mm ± 0.24; bill depth: 3.37 mm ± 0.04; *P* < 0.001) and south-western (mass: 11.35 g ± 0.16; wing chord: 68.09 mm ± 0.30; bill depth: 3.62 mm ± 0.05; *P* < 0.001) populations. North-western populations have smaller tarsi (17.6 mm ± 0.13) and narrower bills (3.94 mm ± 0.06) than south-western (tarsi: 18.25 mm ± 0.16; bill width: 4.61 mm ± 0.08; p < 0.001) and eastern (tarsi: 18.30 mm ± 0.19; bill width: 4.53 mm ± 0.09; p < 0.001) populations. The first canonical analysis examining mass and bill depth explained 77.6% of the variance (Wilks’ lambda = 0.22, χ^2^ = 263.8, *P* < 0.001), and the second discriminant analysis examining differences in tarsus length and bill width accounted for the remaining 22.4% of the variance (Wilks’ lambda = 0.64, χ^2^ = 76.9, *P* < 0.001). Discriminant function analysis assigned 81% of birds to the correct genetic group following cross-validation; 92% of north-western birds and 82.9% of eastern birds were correctly assigned. Among south-western birds, 62.5% of birds were assigned correctly, while 33.9% of south-western birds were assigned to the north-western genetic group.

Differences in morphology were also reported among the subspecies designations (Wilks’ lambda = 0.25, F_12,340_ = 27.5, *P* < 0.001), but rather across all six morphological characters in comparison to the genetic groups (Supporting Information, Table S2). A similar result between the eastern subspecies and eastern genetic group was found: *gilvus* are heavier (14.2 g ± 0.19), have longer wings (69.9 mm ± 0.35), and deeper bills (3.95 mm ± 0.06) than *swainsonii* (mass: 11.6 g ± 0.11; wing chord: 66.9 mm ± 0.20; bill depth: 3.42 mm ± 0.03; *P* < 0.001) and *brewsteri* (mass: 11.3 g ± 0.19; wing chord: 68.9 mm ± 0.35; bill depth: 3.60 mm ± 0.06; *P* < 0.001) populations. The two morphological characters that were diagnostic to the north-western genetic group were also found to be the same in *swainsonii*, with *swainsonii* having smaller tarsi (17.7 mm ± 0.11), and narrower bills (width: 4.06 mm ± 0.06) than both *gilvus* (tarsus: 18.3 mm ± 0.19; width: 4.50 mm ± 0.10; *P* < 0.001) and *brewsteri* (18.4 mm ± 0.19; width: 4.70 mm ± 0.10; *P* < 0.001). One main difference between genetic groups and subspecies comparisons was that the bill length of *brewsteri* is longer (9.74 mm ± 0.22) than both *swainsonii* (8.89 mm ± 0.13; *P* < 0.001) and *gilvus* (9.00 mm ± 0.22; *P* < 0.001). The first canonical analysis examining mass and bill depth explained 81.8% of the variance (Wilks’ lambda = 0.24, χ^2^ = 247.5, *P* < 0.001), and the second discriminant analysis examining differences in tarsus length and bill width accounted for the remaining 18.2% of the variance (Wilks’ lambda = 0.70, χ^2^ = 61.6, *P* < 0.001). Discriminant function analysis assigned 82.7% of birds to the correct genetic group following cross-validation; 90.7% of *swainsonii* individuals and 82.9% of *gilvus* were correctly assigned. Among *brewsteri*, 58.3% of birds were assigned correctly, while 36.1% of *brewsteri* birds were assigned to *swainsonii*.

Overall, our morphological analyses reveal that the eastern genetic group and subspecies *gilvus* differ significantly from the two western genetic groups and subspecies *swainsonii* and *brewsteri*. Among the two western genetic groups and subspecies *swainsonii* and *brewsteri*, there is considerable overlap in both contexts indicating that morphological differences are less pronounced between them.

### Ecological Niche Modelling

Each genetic group’s model from Maxent was evaluated using the area under the curve (AUC) value: north-western AUC = 0.97 (± 0.004), south-western AUC = 0.96 (± 0.003), and eastern AUC = 0.95 (± 0.004). These high AUC values (in proximity to the value of 1), along with test omission curves close to the predicted omission curve, shows that each model performed well. Climatic variables that contributed most were different across each genetic group’s model: eastern (BioClim18 = 35%, BioClim7 = 18%, BioClim11 = 17.5%; precipitation of warmest quarter, annual temperature range, and mean temperature of coldest quarter, respectively), north-western (BioClim14 = 28.8%, BioClim12 = 25.6%, and BioClim11 = 21.5%; precipitation of driest month, annual precipitation, and mean temperature of coldest quarter, respectively), and south-western (BioClim3 = 37.4%, BioClim2 = 22.7%, BioClim1 = 17.5%; isothermality, mean diurnal range, and annual mean temperature, respectively).

Based on our model, the distribution (Fig. 6; bottom) for the eastern genetic group overall resembles the current subspecies boundaries for *gilvus* (Fig. 1), with its range extending west to the Rocky Mountain range. Between the north-western and south-western genetic groups, their predicted distributions in western North America are a little less clear with an abundance of niche overlap (Fig. 6; top and middle). Unlike the cohesion between the eastern genetic group and *gilvus*, the two western genetic groups and the two western subspecies boundaries do not agree based on climatic variables.

**Figure 6.**
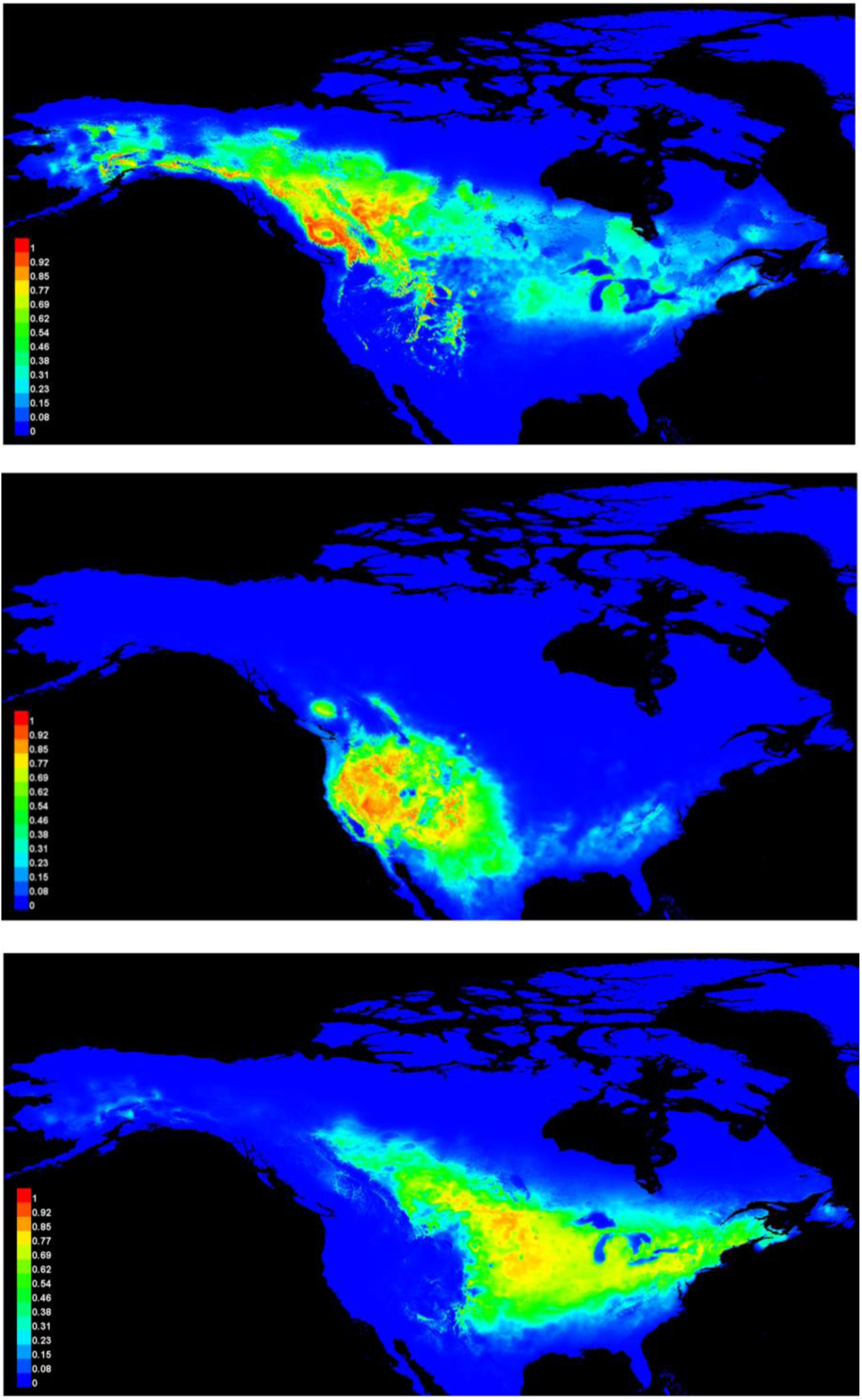
Contemporary ecological niche models for three of the four warbling vireo microsatellite genetic groups (Black Hills are not included): north-western **(top)**, south-western **(middle)**, and eastern **(bottom)**. The logarithmic scale on the left depicts the percent likelihood of habitat suitability based on climate variables.

## DISCUSSION

Taken together, we have conducted the first extensive range-wide population genetic study on the warbling vireo complex to test the congruence of the subspecies designations to genetic data and provide evidence for another potential case of cryptic speciation within the genus *Vireo*. We found a main west – east division for both mtDNA (*cyt b*) and microsatellites, a pattern observed in other widely distributed North American passerines (Milot *et al*., 2000; Clegg *et al*., 2003; Lait & Burg, 2013; Graham *et al*., 2020). The large genetic distances in mtDNA (4.0%) and abrupt split along the Great Plains suggest that the eastern genetic group was isolated in a separate glacial refugium from the two western genetic groups during the Pleistocene (Rising, 1983). Our estimated coalescence time dates 2 Mya, which corresponds to the early Pleistocene (2.5 Mya – 11.5 Kya) (Gibbard *et al*., 2010); however, the timing and role of glaciation in speciation remains controversial (Klicka & Zink, 1997; 1999; Weir & Schluter, 2004; Johnson & Cicero, 2004).

Species in other vireo complexes were found to have split in the early to mid-Pleistocene, including Hutton’s vireo (1.5 Mya) and the solitary vireo (1.3 Mya) (Cicero & Johnson, 1992; 1998). The American Ornithologists’ Union (Banks *et al*., 1997) elevated three subspecies of the solitary vireo complex to species (blue-headed vireo *Vireo solitarius* (Wilson, 1810), Cassin’s vireo *Vireo cassinii* (Xantus de Vesey, 1858) and plumbeous vireo *Vireo plumbeus* (Coues, 1866)) based on *cyt b* sequences (Murray *et al*., 1994) and allozyme variation (Johnson, 1995). Similarly, cryptic populations of eastern and western winter wrens [formerly *Troglodytes troglodytes* (Linnaeus, 1758)] were recognized as two distinct species (winter wren *Troglodytes hiemalis* (Vieillot, 1819) and Pacific wren *Troglodytes pacificus* (Baird, 1864)) following detailed analyses using a range of genetic, acoustic, and morphometric characters (Toews & Irwin, 2008). The genetic distance in *cyt b* between populations of eastern and western warbling vireos in our study (0.040) and previous studies (Murray *et al*., 1994; Lovell *et al*., 2021) is comparable to the distance used when describing species (0.044) rather than subspecies (0.0048) (Nei, 1978; Barrowclough, 1980). Integrated approaches to assess reproductive isolation are increasingly being favored and applied before taxonomic revision. Our general conclusion that the eastern genetic group has experienced prolonged genetic isolation from the two western genetic groups is supported by multiple types of data including mitochondrial and nuclear DNA, morphology, and song.

Within the western populations, the nuclear DNA showed clear separation into two groups, but, morphological and song differences were not as clear, and the two groups observed with this genetic data did not correspond to current distributions of the two western subspecies. The western subspecies morphological characters are described as similar, with overlapping bill and wing morphologies, minor size differences, and dull plumage (Phillips, 1991; Pyle, 1997). MtDNA at both *cyt b* and *ATP* genes revealed a single western lineage with one common haplotype and several haplotypes differing by one to four nucleotide changes. Dividing the western subspecies populations into their respective genetic groups (north-western, south-western, or the Black Hills) provided no additional clarification between the two haplotype networks. Our estimated coalescence times at both mtDNA genes between the north-western and south-western genetic groups, 200,000 – 150,000 years ago, correspond to the last interglacial period or the beginning of the Wisconsin glaciation (Dawson, 1992). Given this recent divergence, the probability of complete lineage sorting to accrue fixed differences in mtDNA and form distinct phylogroups is reduced (Peters *et al*., 2005). Our north-western and south-western genetic groups are distributed north – south, coming into contact in southern Canada and the north-western United States, areas at the southern edge of two ice sheets during the Pleistocene glaciations (Remington, 1968; Swenson & Howard, 2004; 2005). Similar patterns have been observed in other high latitude species (Van Els *et al*., 2012; Chavez *et al*., 2011) and attributed to Pleistocene glacial vicariance.

Our bioacoustic results between both genetic groups and subspecies designations were similar. The eastern genetic group and subspecies *gilvus* produce longer songs with a higher syllable delivery rate, two diagnostic characters from both the western genetic groups and subspecies songs. Between the western subspecies, diagnostic characters in acoustic differences were more muddled, but *swainsonii* songs are shorter and contain fewer syllables than *brewsteri* songs, which reflects our example song spectrograms between the western genetic groups (Fig. 5). A greater proportion of songs were also correctly assigned to each of the western genetic groups rather than to each of the western subspecies, particularly for *brewsteri*. Songs of *gilvus* have been previously described as complex (rhythmic structure; Howes-Jones, 1985) and it has been suggested that complex songs encode species information (Marler, 1960). For example, the complexity at the start of *gilvus* songs were comparable to indigo bunting (*Passerina cyanea*) songs, which gain attention of conspecifics (Shiovitz, 1976; Howes-Jones, 1985). Acoustic divergence has been recognized as a mode of speciation in other cryptic avian species (Irwin *et al*., 2008; Toews & Irwin, 2008), while undifferentiated in others (Garg *et al*., 2016). Song playback experiments in areas of sympatry, or contact zones, would help to resolve the role of singing type as a behavioral and reproductive isolating barrier between eastern and western warbling vireos.

The findings from our morphological analyses were significant in both the subspecies and genetic group comparisons, but the variation in morphology is complex and does not always agree with previous studies. The western subspecies, *brewsteri*, has the longest bill among the three subspecies, which contradicts previous comments of *gilvus* having the longest bill (Phillips, 1991; Lovell, 2010). Minor size differences between *swainsonii* (smaller) and *brewsteri* are also criteria used to define subspecies boundaries (Phillips, 1991), but we found the opposite result with our data; *swainsonii* and the north-western genetic group are slightly heavier (11.59 g and 11.61 g, respectively) than both *brewsteri* and the south-western genetic group (11.25 g and 11.35 g, respectively). While this difference was not statistically significant, it reinforces that morphological differences between the western groups are more subtle. Eastern birds are significantly larger (∼ 2.5 g heavier), have longer wings, and deeper bills than western birds. These differences in size could be attributed to habitat. In Alberta, the eastern birds have been found primarily in open deciduous forest and aspen parkland whereas the western subspecies, *swainsonii*, occurs in mixed forest and boreal forest habitats in the foothills of the Rocky Mountains (Semenchuk, 1992; Lovell *et al*., 2021).

Ecological differences represent barriers for many avian species, including *Phylloscopus* (F. Boie, 1826) warblers (Richman & Price, 1992), *Pycnonotus* (F. Boi, 1826) bulbul species (Lloyd *et al*., 1997), *Empidonax* (Cabanis, 1855) flycatchers (Johnson & Cicero 2002), and other *Vireo* species (Cicero & Johnson, 1998). Grant and Grant (1997) attested that considerable habitat and climate differences maintain species boundaries. Our ecological niche models support a well-defined split between western and eastern genetic groups, corresponding to different precipitation and temperature variables. Several other songbird species pairs with west – east distributions have similar population structure because of local adaptation to refugial temperatures during the Pleistocene glacial cycles (Swenson, 2006), a scenario also observed by Hargrove & Hoffman (2004) in other fauna and flora.

Our study revealed two unique populations: a disjunct western population in the Cypress Hills, located in the south-eastern corner of Alberta and the south-western corner of Saskatchewan, and a population in the Black Hills, located in western South Dakota and north-eastern Wyoming. Based on pollen evidence, both areas were unglaciated during the last glacial maximum and supported boreal forest species (Froiland, 1978; Strong & Hills, 2005). Accompanied by geological processes that created these elevated features among the Great Plains, boreal species became isolated in the Cypress Hills and the Black Hills as the surrounding areas were rapidly replaced with grassland flora with rising temperatures at the start of the Holocene (Strong & Hills, 2005). While both of these locations are embedded within the eastern subspecies range, what little we do know about its preferred habitat (Semenchuk, 1992; Lovell *et al*., 2021) is a plausible explanation for why *gilvus* are not found in these montane areas. Comparisons of allele frequencies from both surrounding south-western (SWCO, CCO, CWY, and NWWY) and eastern (SK, ND, MN, and IL) populations show that genotypes in the Black Hills are more similar to the south-western genetic group, and it is not a hybrid population of the south-western and eastern genetic groups. Additionally, observed heterozygosity between the Black Hills and the south-western and eastern populations does not show a loss in genetic diversity characteristic of a population bottleneck. Disjunct western populations in the Cypress Hills (Chilton, 2003; Metheny *et al*., 2008; Lumley & Sperling, 2011; Dempsey *et al*., 2019) and genetically distinct populations in the Black Hills (Smith, 1963; Weaver *et al*., 2006) have been found across multiple taxonomic groups.

### Conclusions

This study demonstrates that molecular methods are invaluable to bring clarification to taxonomic relationships among taxa with little phenotypic diversity, but also a reminder that each marker (mitochondrial vs nuclear) is limited in the information they provide about evolutionary history. Our main goal was to see if genetic data corroborated subspecies designations in the warbling vireo complex. Secondly, we demonstrated the utility of combining bioacoustic, morphological and ecological niche models to help characterize the variation of cryptic species. We found that overall, our genetic data supported the distribution of the eastern subspecies, *gilvus*, but our microsatellite genetic groups did not support the western subspecies, *swainsonii* and *brewsteri*. More sampling needs to be done to fill the gaps between our north-western and south-western populations to better determine where they come into contact. The warbling vireo complex is comprised of at least two highly divergent genetic groups (eastern and western) that also possess diagnostic acoustic, ecological, and morphological characters in allopatry. Although the observed genetic differentiation between eastern and western warbling vireos is equal to, or greater than, 33 other pairs of North American avian sister taxa that are recognized as separate species (Johnson & Cicero, 2004), it is argued that genetic data alone are weak evidence to imply reproductive isolation (Winker, 2009). Therefore, additional study in western North America of all three genetic groups in areas of sympatry, or contact zones, to evaluate reproductive isolation is proposed before taxonomic revision.

## ACKNOWLEDGMENTS

We thank all of the graduate students and field assistants that helped with field sample collection. A special thanks to the following museums whose voucher sample contributions made this range-wide research possible: the Denver Museum of Nature and Science, the Burke Museum of Natural History and Culture, the Natural History Museum University of Oslo, the University of Michigan Museum, the Field Museum of Natural History, the Cleveland Museum of Natural History, and the Royal Alberta Museum. We would like to acknowledge the work of the Cornell Lab of Ornithology, and all of its collaborators, in the eBird database. We also thank the editor and the three anonymous reviewers for their comments that helped improve the manuscript.

Samples were collected under the appropriate provincial, state, and federal permits. This work was funded by the Natural Sciences and Engineering Research Council of Canada (NSERC Discovery Grant RGPIN-2019-05068) (T.M.B.) and the National Science Foundation (DEB0814841) (G.M.S). The authors declare they have no conflicts of interest.

## DATA AVAILABILITY

The sequence data underlying this article are available in GenBank with the following accession numbers: MZ020223 - MZ020331.

## SUPPORTING INFORMATION

**Table S1.**
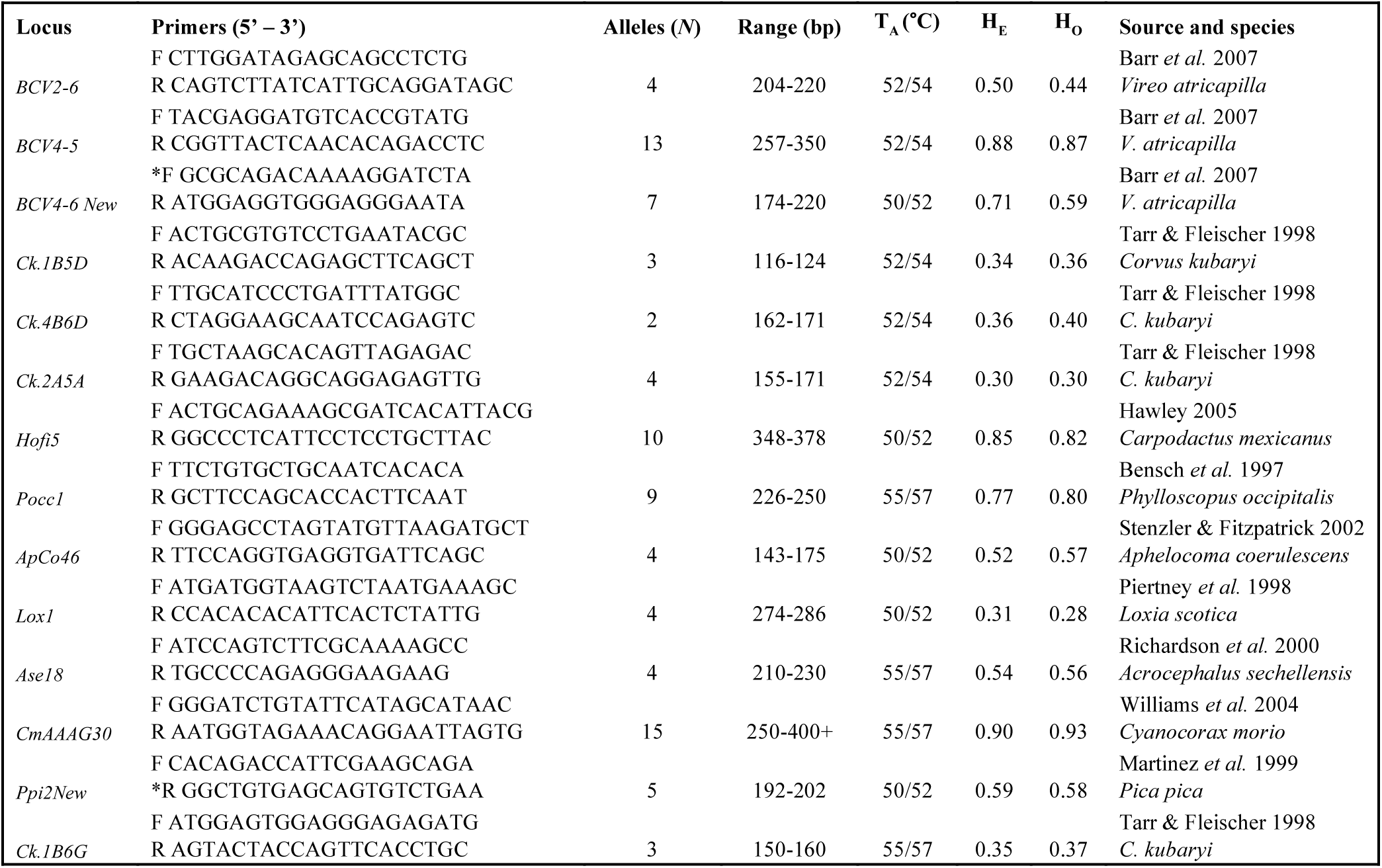
Microsatellite primer names and sequences, average number of alleles per locus (*N*), size range (bp), annealing temperature (T_A_), average expected and observed heterozygosity (H_E_/H_O_), and source with the species the primers were originally designed for. Sequences marked with an asterisk were modified from the original source.

**Table S2.** Values for mean, standard error (SE) and 95% confidence intervals (CI) lower bound (LB) and upper bound (UB) from the morphological comparisons among subspecies and genetic groups. Significant comparisons (Sig) are bolded.

| Variable | Subspecies | Mean | SE | Sig | 95% CI LB | 95% CI UB | Genetic Group | Mean | SE | Sig | 95% CI LB | 95% CI UB |
| --- | --- | --- | --- | --- | --- | --- | --- | --- | --- | --- | --- | --- |
| Mass (g) | *swainsonii* | 11.59 | ± 0.11 | a | 11.38 | 11.81 | North-western | 11.61 | ± 0.13 | a | 11.35 | 11.86 |
|  | *brewsteri* | 11.25 | ± 0.19 | a | 10.87 | 11.63 | South-western | 11.35 | ± 0.16 | a | 11.04 | 11.67 |
|  | *gilvus* | 14.16 | ± 0.19 | **b** | 13.79 | 14.54 | Eastern | 14.16 | ± 0.19 | **b** | 13.78 | 14.54 |
| Wing Chord<br>(mm) | *swainsonii* | 66.98 | ± 0.20 | **a** | 66.59 | 67.38 | North-western | 67.04 | ± 0.24 | **a** | 66.57 | 67.52 |
|  | *brewsteri* | 68.85 | ± 0.35 | **b** | 68.17 | 69.54 | South-western | 68.09 | ± 0.30 | **b** | 67.49 | 68.68 |
|  | *gilvus* | 69.96 | ± 0.35 | **c** | 69.27 | 70.64 | Eastern | 69.96 | ± 0.36 | **c** | 69.24 | 70.67 |
| Tarsus<br>(mm) | *swainsonii* | 17.68 | ± 0.11 | **a** | 17.46 | 17.89 | North-western | 17.60 | ± 0.13 | **a** | 17.35 | 17.85 |
|  | *brewsteri* | 18.40 | ± 0.19 | b | 18.03 | 18.78 | South-western | 18.26 | ± 0.16 | b | 17.94 | 18.57 |
|  | *gilvus* | 18.29 | ± 0.19 | b | 17.92 | 18.66 | Eastern | 18.30 | ± 0.19 | b | 17.93 | 18.68 |
| Bill Length<br>(mm) | *swainsonii* | 8.89 | ± 0.13 | a | 8.63 | 9.15 | North-western | 8.93 | ± 0.15 | a | 8.63 | 9.23 |
|  | *brewsteri* | 9.75 | ± 0.23 | **b** | 9.30 | 10.19 | South-western | 9.38 | ± 0.19 | a | 9.00 | 9.76 |
|  | *gilvus* | 9.00 | ± 0.23 | a | 8.56 | 9.45 | Eastern | 9.00 | ± 0.23 | a | 8.55 | 9.46 |
| Bill Depth<br>(mm) | *swainsonii* | 3.42 | ± 0.03 | **a** | 3.36 | 3.49 | North-western | 3.37 | ± 0.04 | **a** | 3.30 | 3.44 |
|  | *brewsteri* | 3.60 | ± 0.06 | **b** | 3.50 | 3.71 | South-western | 3.62 | ± 0.05 | **b** | 3.53 | 3.71 |
|  | *gilvus* | 3.95 | ± 0.06 | **c** | 3.84 | 4.06 | Eastern | 3.96 | ± 0.05 | **c** | 3.85 | 4.07 |
| Bill Width<br>(mm) | *swainsonii* | 4.06 | ± 0.06 | **a** | 3.95 | 4.17 | North-western | 3.94 | ± 0.06 | **a** | 3.82 | 4.07 |
|  | *brewsteri* | 4.66 | ± 0.10 | b | 4.47 | 4.85 | South-western | 4.61 | ± 0.08 | b | 4.46 | 4.77 |
|  | *gilvus* | 4.51 | ± 0.10 | b | 4.32 | 4.70 | Eastern | 4.53 | ± 0.09 | b | 4.35 | 4.72 |

